# B cell c-Maf signaling promotes tumor progression in animal models of pancreatic cancer and melanoma

**DOI:** 10.1101/2024.09.30.615831

**Authors:** Qian Zhong, Hongying Hao, Shu Li, Yongling Ning, Hong Li, Xiaoling Hu, Kelly M. McMasters, Jun Yan, Chuanlin Ding

## Abstract

**Background:** The role of B cells in anti-tumor immunity remains controversial, with studies suggesting the pro-tumor and anti-tumor activity. This controversy may be due to the heterogeneity in B cell populations, as the balance among the subtypes may impact tumor progression. The immunosuppressive regulatory B cells (Breg) release IL-10 but only represent a minor population. Additionally, tumor-specific antibodies (Ab) also exhibit anti-tumor and pro-tumor function dependent on the Ab isotype. Transcription factor c-Maf has been suggested to contribute to the regulation of IL-10 in Breg, but the role of B cell c-Maf signaling in anti-tumor immunity and regulating antibody responses remain unknown.

**Methods:** Conditional B cell c-Maf knockout (KO) and control mice were used to establish a KPC pancreatic cancer model and B16.F10 melanoma model. Tumor progression was evaluated. B cell and T cell phenotypes were determined by flow cytometry, mass cytometry, and cytokine/chemokine profiling. Differentially expressed genes in B cells were examined by using RNA-seq. Peripheral blood samples were collected from healthy donors and melanoma patients for B cell phenotyping.

**Results:** Compared to B cells from spleen and lymph nodes, B cells in the pancreas exhibited significantly less follicular phenotype and higher IL-10 production in naïve mice. c-Maf deficiency resulted in a significant reduction of CD9^+^ IL-10-producing Breg in the pancreas. PDAC progression resulted in accumulation of circulating B cells with follicular phenotype and less IL-10 production in the pancreas. Notably, B cell c-Maf deficiency delayed PDAC tumor progression and resulted in pro-inflammatory B cells. Further, tumor volume reduction and increased effective T cells in the tumor-draining lymph node (TDLN) were observed in B cell c-Maf KO mice in the B16.F10 melanoma model. RNA-seq analysis of isolated B cells revealed that B cell c-Maf signaling modulates immunoglobulin (Ig)-associated genes and tumor specific antibody production. We furthermore demonstrated c-Maf-positive B cell subsets and increase of IL-10-producing B cells after incubation with IL-4 and CD40L in the peripheral blood of melanoma patients.

**Conclusion:** Our study highlights that B cell c-Maf signaling drives tumor progression through the modulation of Breg, inflammatory responses, and tumor-specific Ab responses.

**What is already known on this topic:** The net effect of B cells on tumor immunity depends on the balance of various B cell subtypes. c-Maf has been suggested to contribute to the regulation of IL-10 in regulatory B cells (Breg), but the role of B cell c-Maf signaling in anti-tumor immunity remains unknown.

**What this study adds:** This study shown that B cell c-Maf signaling drives tumor progression in pancreatic cancer and melanoma. We defined different anti-tumor mechanisms of B cell c-Maf deficiency in two tumor models. Specifically, c-Maf signaling modulates the pro-inflammatory phenotype of B cells in the KPC tumor-bearing pancreas and tumor-specific antibody responses in tumor draining lymph nodes (TDLN) of melanoma.

**How this study might affect research, practice or policy:** These studies indicate that inhibition of c-Maf signaling is a novel and promising approach for immunotherapy in pancreatic cancer and melanoma.

## BACKGROUND

B cells are known to regulate immunity by producing specific antibodies, secreting cytokines, serving as antigen-presenting cells to promote T cell responses, and forming tumor-associated tertiary lymphoid structures (TLSs). Previous studies have demonstrated that B cells are significantly expanded in the pancreas and contribute to the development of pancreatic ductal adenocarcinoma (PDAC) (1, 2). Further, tumor progression results in significant B cell expansion and activation in tumor draining lymph nodes (TDLN) (3, 4), which can simultaneously modulate anti-tumor immunity and metastatic potential (5, 6).

Although B cells are generally thought to augment immune responses and anti-tumor immunity (7–11), B cells also suppress immune responses in a variety of mouse models of cancer (12–15). This controversy may be due to the fact that B cells have multiple subsets depending on their stage of development and function. Regulatory B cells (Breg) exhibit immunosuppressive functions by secreting cytokines (IL-10, IL-35, and TGF-β) and inducing regulatory T cell differentiation (2, 15, 18–20), but the immunosuppressive Breg represent a minor cell population of B cells. The pro-tumor effect of Breg might be overcome by the presence of other subsets of B cells. Therefore, the net effect of B cells on tumor immunity depends on the balance of various B cell subtypes (16, 17). In addition, the transcriptional regulation of Breg in the tumor microenvironment and TDLN remains poorly understood. We postulate that targeting of Breg could improve anti-tumor responses by reversing their pro-tumor effects.

It has been shown that transcription factor c-Maf plays essential roles in regulating of T cells and myeloid cells (18). Our previous studies have demonstrated that c-Maf is overexpressed in tumor-associated macrophages and γδT17 cells and is associated with their pro-tumor effect (19, 20). Although c-Maf has been suggested to contribute into Breg differentiation (21–23), the roles of B cell c-Maf signaling in anti-tumor immunity remain unknown. In this study, we demonstrate that specific knockout of c-Maf in B cells inhibits tumor progression in animal models of PDAC and melanoma, which is associated with an increase of effector T cells. In addition to regulation of IL-10 production, our study also suggests that c-Maf can regulate B cell inflammatory responses and tumor-specific Ab responses in mouse models of PDAC and melanoma, respectively.

## RESULTS

### c-Maf regulates pancreas B1 B cells and B cell IL-10 expression

To investigate the roles of c-Maf in regulating B cell development and function, we analyzed the data from The Immunological Genome Project (http://www.immgen.org) and found that c-Maf is expressed at very low levels in B cells. This finding was confirmed by Flow cytometry analysis. However, we found that c-Maf is detectable in CD9^+^ B cells which is IL-10-producing B cell subset (24). Western blotting verified that c-Maf is remarkably downregulated in CD9^+^ B cells of conditional B cell c-Maf knockout (KO) mice (online supplemental figure 1A and B). We further used IL-10^gfp^ reporter mice and found that c-Maf is expressed in IL-10^+^ B cells at relative higher levels (online supplemental figure 1C).

Prior studies highlight that tumor location contributes into the discrepancy of B cell function in tumors (8, 25, 26). Naïve peripheral B cells can be generally divided into three subsets: follicular B cells, marginal zone (MZ) B cells and B1 B cells. The comparison of B cell subsets among tissues and the impact of B cell c-Maf signaling have not been reported. To better understand roles of B cell in tumor immunity in different tissues, we started by examining the distribution of B cell subpopulations in spleen, lymph nodes (LN), and pancreas from naive control and B cell c-Maf KO mice using Flow cytometry, and the gating strategy was shown in supplemental figure 2. As shown in figure 1A, B cells in the pancreas exhibited significantly less follicular phenotype compared to B cells from the spleen and LN. c-Maf deficiency did not alter the composition of follicular B cells and MZ B cells. Naïve B cells express B cell receptor (BCR), including IgM and IgD isotype. Compared to the B cells in spleen and LN, pancreas had more IgD^low^IgM^+^ B cells and less IgD^+^IgM^low^ B cells. c-Maf deficiency did not impact the composition of IgD^+^IgM^low^ B cells and IgD^low^IgM^+^ B cells in these tissues (figure 1B).

**Figure 1.**
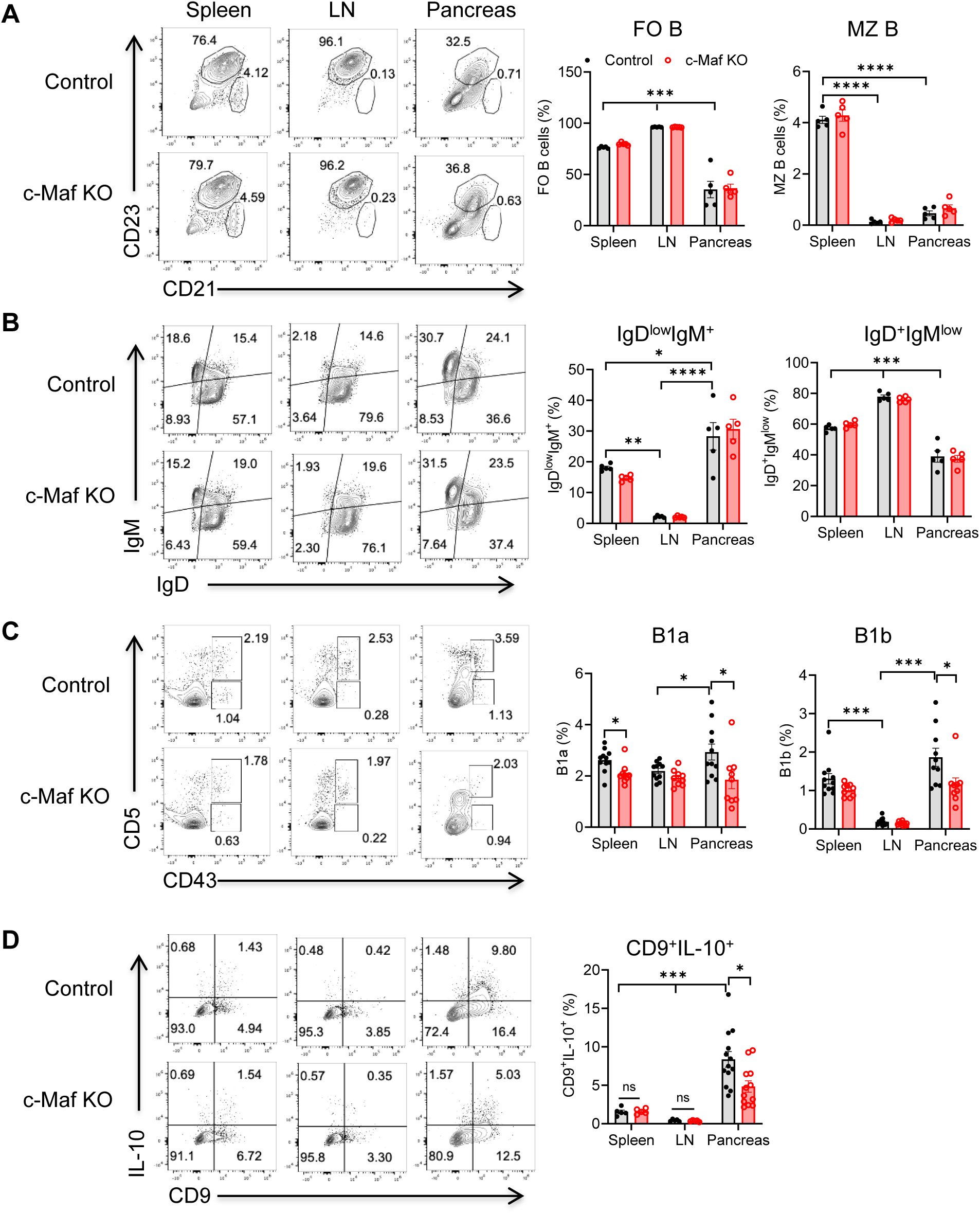
B cell subsets and B cell IL-10 expression in spleen, lymph node, and pancreas of naïve control and c-Maf KO mice. (A) Follicular B cells (FO, CD21^low^CD23^+^) and marginal zone B cells (MZ, CD21^+^CD23^−^) within CD19^+^ B cells. (B) IgD and IgM expression by B cells. (C) B1a (CD43^+^CD5^+^) and B1b (CD43^+^CD5^−^) within CD19^+^IgM^+^ B cells. (D) IL-10 production by CD9^+^ B cells after 4-6 hour of PMA/Ionomycin/LPS stimulation. Representative plots and summarized results are shown. Each point represents an individual mouse (n=5-13). *p<0.05; **p<0.01; ***p<0.001; ****p<0.0001. IL-10, interleukin 10; PMA, phorbol 12-myristate 13-acetate; LPS, lipopolysaccharides; FO: follicular B cells; MZ: marginal zone B cells.

B1 B cells are a small but unique subpopulation of B cells that can constitutively produce IL-10 (27). There are two subsets of B1 B cells: CD5^+^ B1a and CD5^−^ B1b. Compared to the spleen and LN, the pancreas was infiltrated with more B1a and B1b cells. c-Maf deficiency decreased pancreas B1a and B1b cells as well as spleen B1a cells. No significant changes of B1a and B1b cells were observed in LN between control and B cell c-Maf KO mice (figure 1C). We further examined IL-10 production in spleen, LN, and pancreas B cells from control and B cell c-Maf KO mice. Notably, we observed a significant greater population of CD9^+^IL-10^+^ B cells in the pancreas, which was correlated with an increase of B1 B cells in the pancreas. c-Maf deficiency significantly reduced the proportion of CD9^+^ IL-10-producing B cells in the pancreas (figure 1D). Together, these data suggest that B cells in the pancreas exhibit a distinct phenotype. B cell c-Maf signal modulates B cell IL-10 expression in the pancreas under steady state condition.

### Tumor progression induces complexity of B cell phenotype in pancreas

Previous studies have shown that B cells exhibit immunosuppressive roles in the KPC mouse model of pancreatic cancer, especially IL-35-producing B cells (14, 15, 17). However, it was also reported that B cells upregulate several proinflammatory and immunostimulatory genes upon tumor progression (28). Therefore, the phenotype and function of B cells as a whole population in the KPC tumor model are still not well understood. Subsets of infiltrating B cells in the pancreas during tumor progression have not been thoroughly investigated. To understand the changes of B cell phenotype and function during tumor progression, we evaluated the distribution of B cells and their subpopulations in the pancreas from tumor-free and KPC tumor-bearing mice by flow cytometry. KPC tumor development induced a significant infiltration of CD19^+^ B cells in the pancreas, including increased percentage of CD19^+^ B cells within CD45^+^ leukocytes (figure 2A) and increased absolute number of B cells (figure 2B). The expanded B cells mainly exhibited follicular B cell phenotype (CD23^+^CD21^low^) (figure 2C). Accordingly, IgD^+^IgM^low^ B cells were increased whereas IgD^low^IgM^+^ B cells were decreased in the tumor-bearing pancreas (figure 2D).

**Figure 2.**
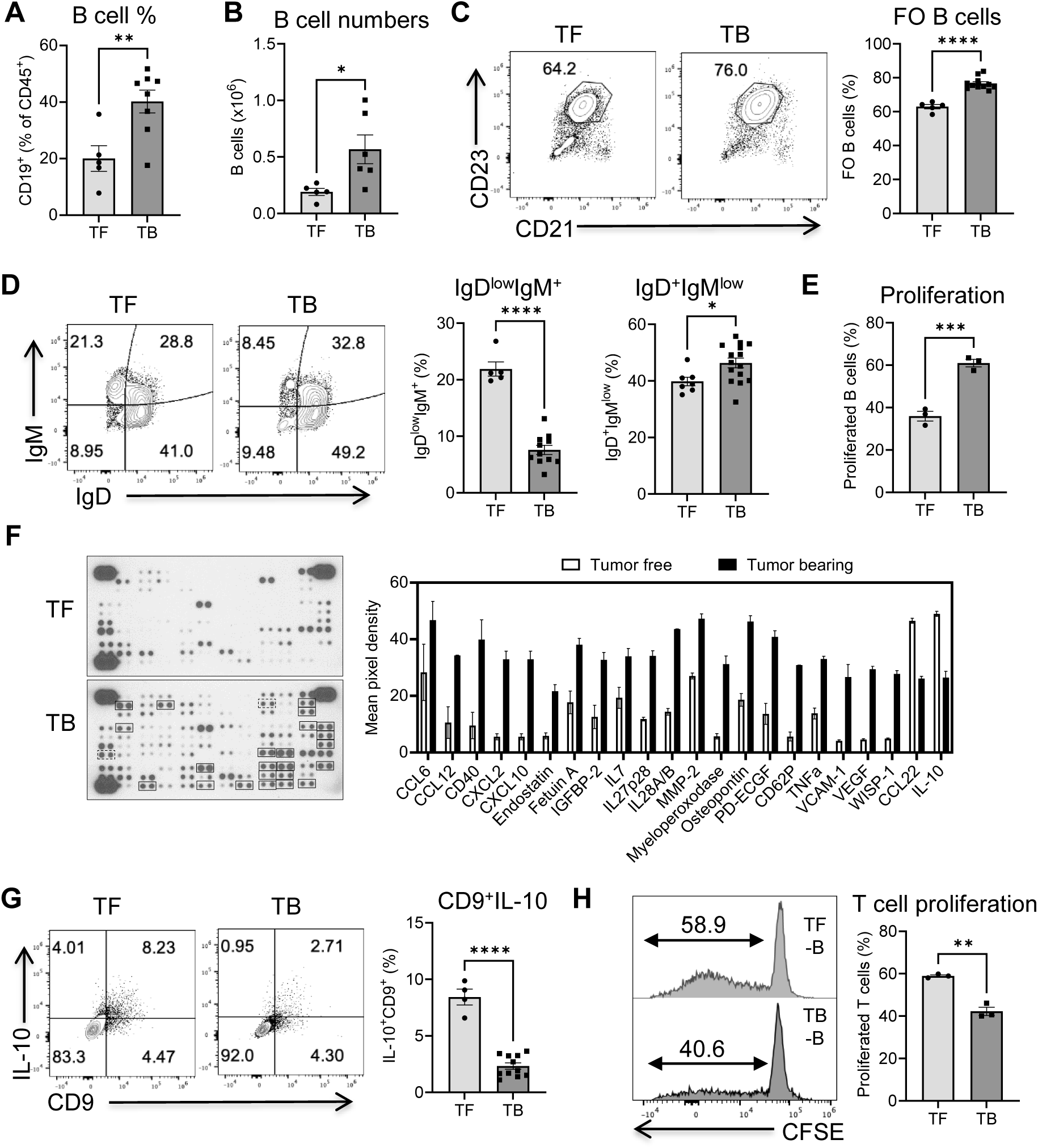
Tumor progression induces a complex B cell phenotype in pancreas. 8-10 weeks old B6 wild-type mice were implanted orthotopically with KPC pancreatic cancer cells. (A) Percentages of CD19^+^ B cells within CD45^+^ leukocytes in tumor-free (TF) and tumor-bearing (TB) mice. (B) Absolute cell numbers of B cells in the pancreas of TF and TB mice. (C) Percentages of follicular B cells within CD19^+^ B cells. (D) IgD and IgM expression by B cells. Each point represents an individual mouse (n=5-11). (E) B cell proliferation after 3 days culture in the presence of LPS (1 μg/ml). (F) Cytokine/chemokine array of culture supernatants of isolated B cells. The mean pixel intensity of differentially expressed molecules was measured using ImageJ software. (G) IL-10 production by B cells after *ex vivo* stimulation with PMA/Ionomycin/LPS. (H) T cell proliferation of OT-II CD4^+^ T cells after co-culture with isolated B cells in the presence of OVA for 4 days. *p<0.05; **p<0.01; ***p<0.001; ****p<0.0001. TF, tumor-free; TB, tumor-bearing.

To examine B cell phenotypic changes upon tumor development, B cells were isolated from tumor-free and KPC tumor-bearing pancreas and stimulated with LPS for 3 days. As shown in figure 2E, the tumor-infiltrating B cells proliferated extensively upon LPS stimulation. Further, isolated B cells were stimulated with PMA/Ionomycin/LPS for 24 hours and culture supernatants from stimulated B cells were collected for cytokine and chemokine assay. The array analysis revealed that tumor-infiltrating B cells secreted a vast array of proteins upon *ex vivo* stimulation. Several cytokines and chemokines, such as CCL6, CCL12, CXCL2, CXCL10, IL7, IL27, IL28, and TNF-α, were highly expressed in tumor-infiltrating B cells. Interestingly, down-regulation of IL-10 and CCL22 was observed in the culture supernatants of tumor-infiltrating B cells (figure 2F). In agreement, the percentage of CD9^+^IL-10^+^ B cells was also decreased in the tumor-bearing pancreas (figure 2G). This could be explained by the fact that more follicular B cells are circuited and infiltrated into pancreas in response to tumor progression.

To determine net impact of B cell on T cell activation, total B cells (CD45^+^CD19^+^ B cells) were sorted from the pancreas of tumor-free and KPC tumor-bearing mice and co-cultured with purified CD3^+^CD4^+^ OT-II T cells. As expected, B cells from tumor-bearing mice exhibited decreased antigen presentation function for T cell activation (figure 2H). Together, these data illustrate that tumor progression induces a complex B cell phenotype in the pancreas. B cells as a whole population exhibit a proinflammatory phenotype and decreased antigen-presenting capability.

### B cell c-Maf deficiency delays tumor progression in mouse PDAC model

Given that c-Maf regulates B cell IL-10 expression in pancreas, we investigated the roles of B cell intrinsic c-Maf signaling in regulating anti-tumor immunity in mouse PDAC model. 1×10^5^ KPC-GFP cells were mixed with Matrigel at 1:1 ratio, and 50 μl mixture were orthotopically implanted into the tail of the pancreas of 8-10 weeks old B cell c-Maf KO and control mice. Pancreas were collected at day 18-21 post tumor cell implantation. As shown in figure 3A, both pancreas weight and the amount of GFP^+^ KPC tumor cells showed a decrease in B cell c-Maf KO mice. IL-10^+^ B cells were also decreased in c-Maf KO mice, although overall B cell percentages were not changed (figure 3A). Accordingly, effective T cells (CD4^+^IFN-γ^+^, CD8^+^IFN-γ^+^) were significantly increased in the tumor-bearing pancreas of B cell c-Maf KO mice (figure 3B). To determine whether B cell c-Maf deficiency directly impacts T cell activation, total CD19^+^ B cells were sorted from KPC tumor-bearing pancreas of control and c-Maf KO mice and co-cultured with FACS-isolated CD3^+^CD4^+^ OT-II T cells in the presence of antigen ovalbumin (OVA). T cells proliferated at similar rate when co-cultured with control or c-Maf-deficient B cells (figure 3C), suggesting that B cell c-Maf deficiency has no direct effect on T cell activation when using the whole B cell population as antigen-presenting cells (APCs).

**Figure 3.**
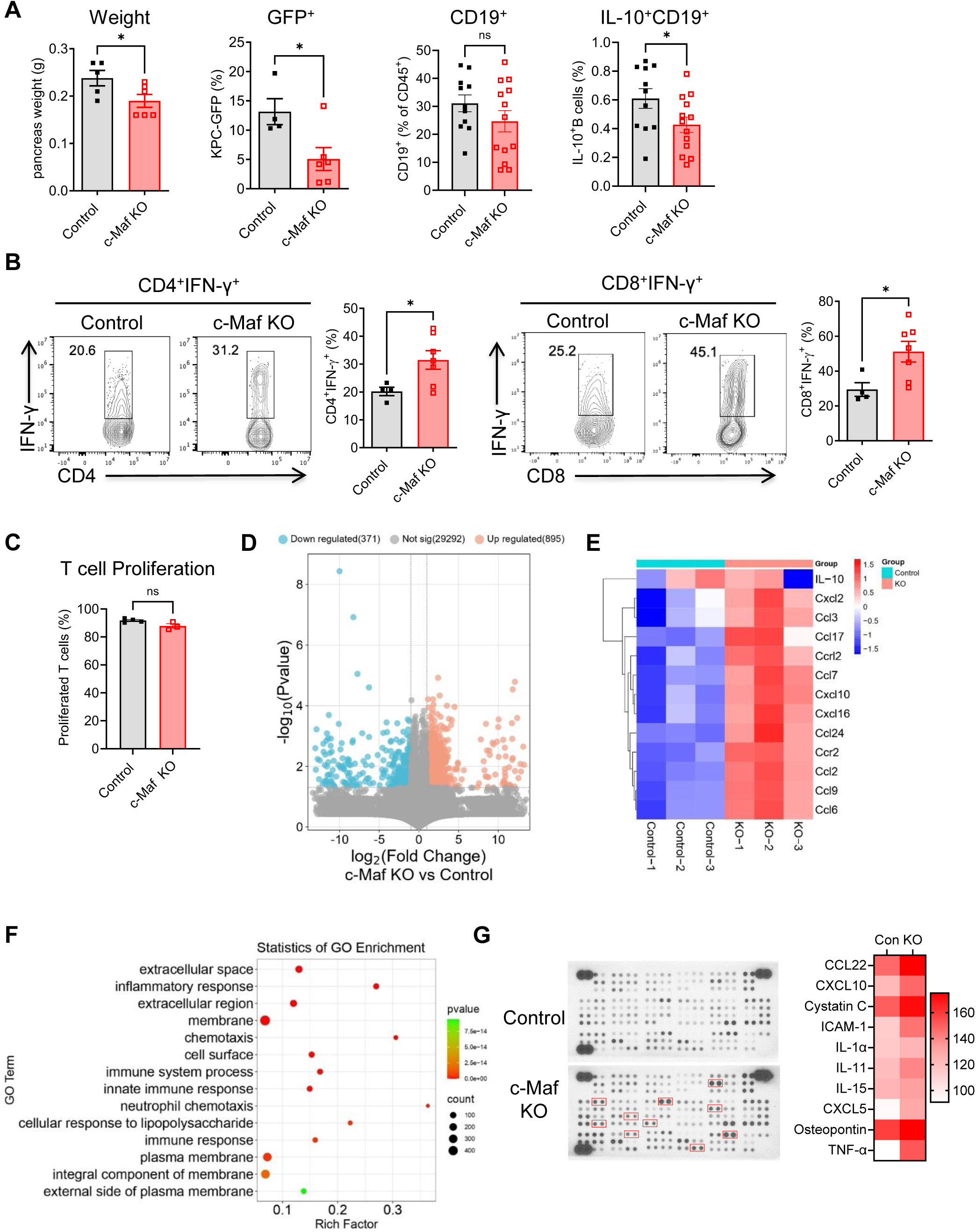
B cell c-Maf deficiency delays tumor progression in the PDAC mouse model. (A) Pancreas weight, percentages of GFP^+^ tumor cells within CD45^−^ pancreas cells, CD19^+^ B cells within CD45^+^ leukocytes, and IL-10-producing B cells of KPC tumor-bearing control and c-Maf KO mice. Each point represents an individual mouse (n=5-13). (B) IFN-γ production by CD4^+^ and CD8^+^ T cells after ex vivo restimulation with PMA/Ionomycin. (C) T cell proliferation of OT-II CD4^+^ T cells after co-culture with FACS-isolated control and c-Maf KO B cells. (D) Volcano plot showing fold change (FC) and *p* value for the comparison of B cells from control versus c-Maf KO mice based on RNA-seq data (n = 3). (E) Heatmap showing the clustering of chemokines in two groups based on log-relative abundances. (F) GO enrichment analysis of genes defining the 14 most significant terms. (G) Cytokine/chemokine array of culture supernatants of isolated B cells. The mean pixel intensity of differentially expressed molecules was measured using ImageJ software. *p<0.05; ns, not significant.

To investigate how c-Maf deficiency in B cells affects tumor progression, we analyzed gene profiles of B cells using RNA-seq. Altogether, 1,266 genes were differentially expressed between tumor-educated control and c-Maf-deficient B cells (figure 3D). Interestingly, several chemokines, including CCL3, CCL6, CCL9, CCL7, CCL17, CXCL12, CXCL10, and CXCL16, were upregulated in c-Maf-deficient B cells (figure 3E). Next, functional annotation of genes in tumor educated c-Maf deficient B cells compared with control B cells was performed. c-Maf deficient B cells were uniquely enriched in gene sets associated with inflammatory response and chemotaxis (figure 3F). We further isolated B cells and stimulated with PMA/Ionomycin/LPS for 24 hours and collected culture supernatants for cytokine and chemokine analysis. Compared to the B cells from tumor-bearing control mice, B cell c-Maf deficiency resulted in an increase of CCL22, Cysatin C, CXCL5, CXCL10, IL-11, and TNF-α (figure 3G). These molecules have been shown to have pro-inflammatory effects in previous studies (29–31). These data are consistent with previous findings that B cells can sustain inflammation and predict response to immune checkpoint blockade (32), and that pro-inflammatory chemokines can boost NK and T cell recruitment leading immunological tumor control (33). Together, our data suggest that B cell c-Maf signaling exhibits a pro-tumor effect in the KPC tumor model via regulation of a pro-inflammatory phenotype in B cells.

### B cell c-Maf deficiency reduces B16.F10 melanoma tumor progression

For many solid tumors, lymph node remodeling contributes into both metastatic potential and immune surveillance (5). Previous studies have shown that B cells are significantly expanded in the tumor-draining lymph node (TDLN) and promote breast cancer metastasis (3). It was also reported that there is accumulation of regulatory transitional 2-marginal zone precursor (T2-MZP) B cells in TDLN of B16.F10 melanoma tumor-bearing mice (4). Our data confirmed the expansion of B cells in the TDLN of B16.F10 tumor (figure 4A). Importantly, IL-10-producing B cells were also increased in the TDLN of B16.F10 tumor (figure 4B). To determine whether B cell c-Maf signaling plays roles in melanoma, control and B cell c-Maf KO mice were injected subcutaneously (s.c.) with B16.F10 melanoma cells in the flank. As shown in figure 4C, B16.F10 tumor progression in B cell c-Maf KO mice was significantly reduced compared to control mice.

**Figure 4.**
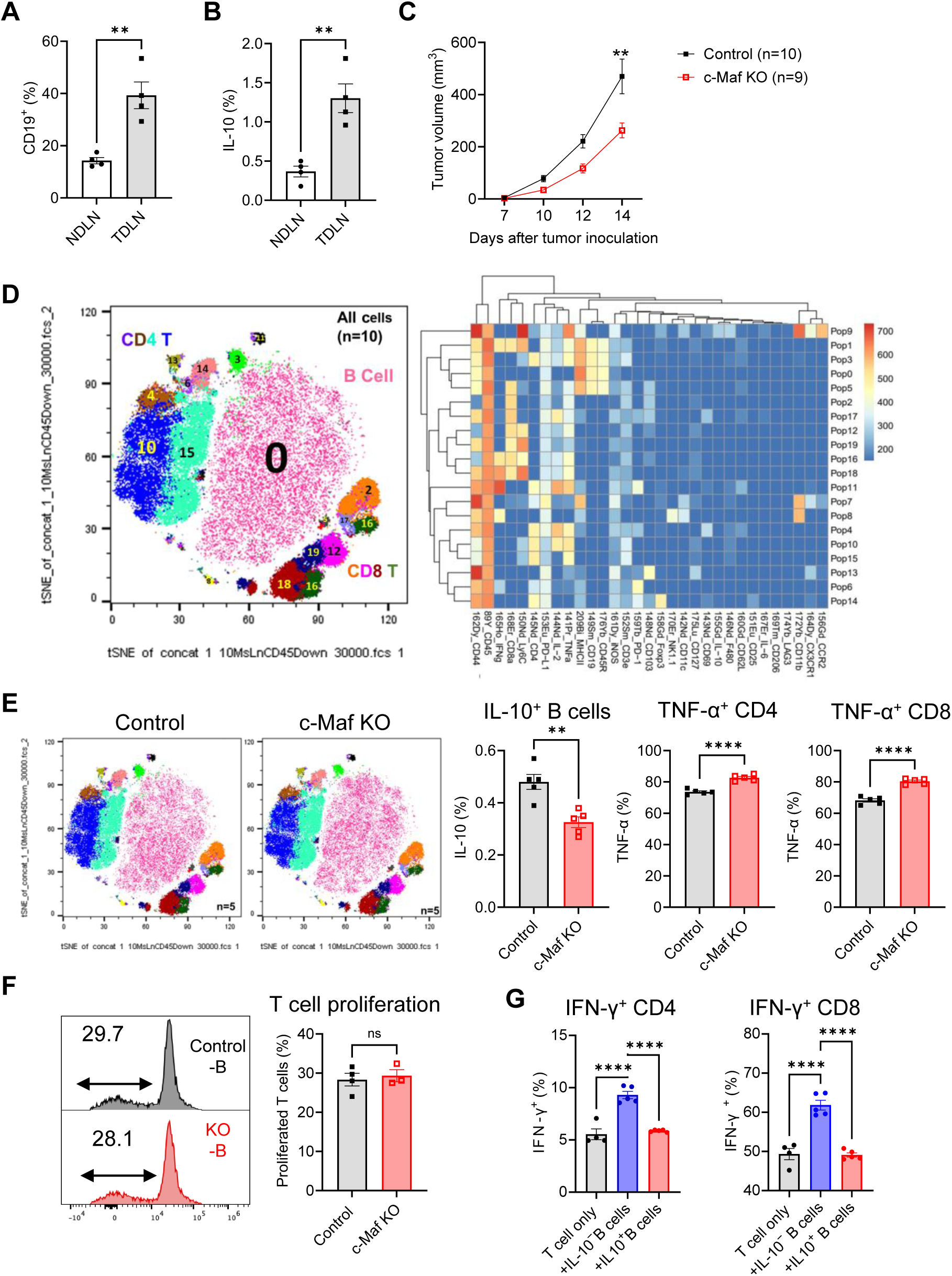
B cell c-Maf deficiency inhibits B16.F10 melanoma tumor progression. (A) Percentages of CD19^+^ B cells within CD45^+^ leukocytes in TDLN of tumor-bearing mice. (B) Percentages of IL-10-producing B cells in TDLN of tumor-bearing mice compared to that in NDLN. (C) Tumor progression in control and c-Maf KO mice (n=9-10). (D) viSNE analysis of CyTOF immunophenotyping of TDLN from tumor-bearing control and c-Maf KO mice, all samples combined (n=10, left). FlowSOM clustering into 20 final immune cell types showing as a normalized expression heatmap (right). (E) viSNE analysis of CyTOF immunophenotyping in TDLN of control and c-Maf KO mice (n=5). Percentages of IL-10-producing B cells, TNF-α^+^ CD4^+^ and CD8^+^ T cells were summarized. (F) T cell proliferation of OT-II CD4^+^ T cells after co-culture with isolated B cells (G) IFN-γ production by anti-CD3 mAb-activated CD4^+^ and CD8^+^ T cells after co-culture with IL-10^−^ and IL-10^+^ B cells. **p<0.01; ****p<0.0001; ns, not significant. NDLN, non-draining lymph nodes; TDLN, tumor-draining lymph nodes.

Next, we used mass cytometry (CyTOF) to determine the effect of B cell c-Maf deficiency on immune cell profiling within TDLN after *ex vivo* re-stimulation with PMA/Ionomycin plus LPS in the presence of brefeldin A for 4-6 hours. FlowSOM analysis identified a total of 20 immune cell subtypes, including B cells (population 0, 1, 3, and 5), CD4 T cells (population 4, 10, 15, 6, 14, and 11), and CD8 T cells (population 2,12, 16, 17, 18, and 19) (figure 4D). Although B c-Maf deficiency did not impact the composition of these major cell subsets, IL-10 production was reduced in c-Maf-deficient B cells, which was associated with an increase of effective TNF-α^+^ CD4^+^ and CD8^+^ T cells (figure 4E).

To evaluate whether B cell c-Maf deficiency has direct effect on T cell activation, total CD19^+^ B cells were sorted from TDLN of control and c-Maf KO B16.F10 tumor-bearing mice and co-cultured with FACS-isolated CD3^+^CD4^+^ OT-II T cells in the presence of antigen ovalbumin (OVA). T cells proliferated at similar rate when co-cultured with control or c-Maf-deficient B cells (figure 4F), suggesting that B cell c-Maf deficiency has no direct effect on T cell activation when using the whole B cell population as APCs. This could be explained by the fact that immunosuppressive B cells represent a relatively minor cell population. To confirm this possibility, we cultured B cells from IL-10^gfp^ reporter mice using NIH3T3/CD40LB feeder cells and cytokines (IL-4, IL-21) (34). IL-10-producing B cells (eGFP^+^) and IL-10-negative B cells (eGFP^−^) were sorted and co-cultured with anti-CD3 activated T cells (figure 4G and online supplemental figure 3), IL-10-negative B cells enhanced IFN-γ production by CD4 and CD8 T cells, whereas IL-10-producing B cells exhibited immunosuppressive effects compared to IL-10-negative B cells. Together, these data suggest that B cell c-Maf signaling exhibits pro-tumor effects in melanoma, which might not be through direct B cell/T cell interaction.

### c-Maf regulates immunoglobulin related genes in B cells and tumor-specific antibody production

To understand how B cell c-Maf signaling regulates anti-tumor immunity in melanoma, we purified B cells from LN of naïve mice and TDLN of B16.F10 tumor-bearing control and B cell c-Maf KO mice and performed RNA-seq. A comparison of differentially expressed genes (DEGs) among different groups was illustrated in a Venn diagram (figure 5A). Several genes related to immunoglobulin, such as *Jchain, Ighg2b, Ighv1-37, Igkv3-12, Igkv8-21, Iglv1, Ighg3*, were significantly upregulated in the tumor-bearing B cells compared to the naïve B cells (figure 5B and 5F). B cell c-Maf deficiency resulted in a significant decrease of these genes in tumor-bearing animals (figure 5C and 5F). Gene ontology enrichment analysis showed that factors involved in circulating immunoglobulin complex, immunoglobulin production, and external side of plasma membrane were enriched in the tumor-bearing B cells compared to naïve B cells (figure 5D). These factors were regulated by c-Maf because these pathways were significantly enriched in the c-Maf deficient tumor-bearing B cells compared to control B cells (figure 5E), suggesting important roles of c-Maf in regulating Ab responses.

**Figure 5.**
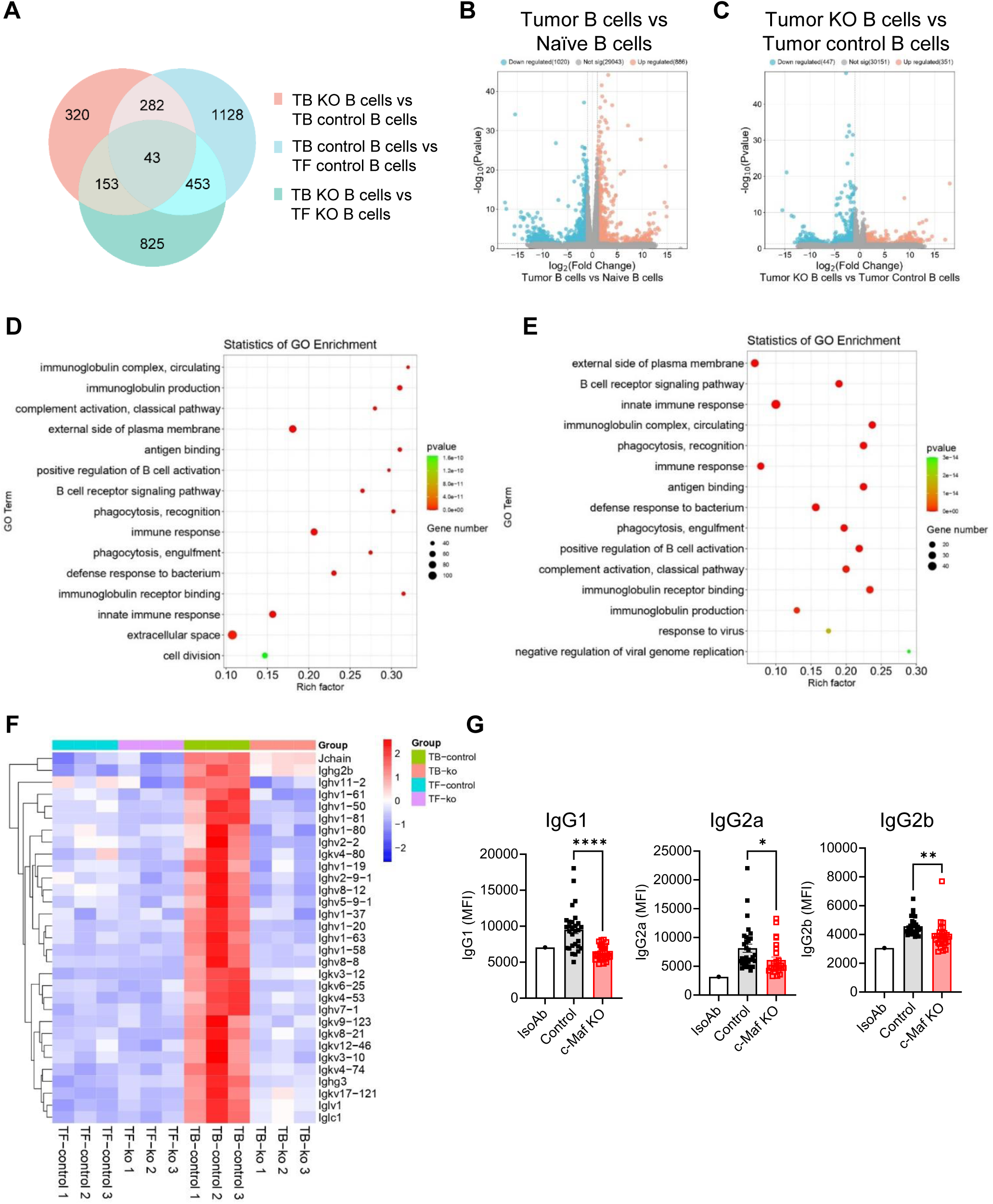
c-Maf regulates immunoglobulin-related genes in B cells and tumor-specific antibody production. (A) Venn diagram showing common and unique genes among the three comparisons. (B) Volcano plot showing fold change (FC) and *p* value for the comparison of B cells from tumor-bearing versus tumor-free naïve mice (n = 3). (C) Volcano plot showing fold change (FC) and *p* value for the comparison of B cells from tumor-bearing c-Maf KO versus control mice (n = 3). (D) GO enrichment analysis of genes defining the 15 most significant terms in B cells from tumor-bearing versus tumor-free naïve mice. (E) GO enrichment analysis of genes defining the 15 most significant terms in B cells from tumor-bearing c-Maf KO versus control mice. (F) Heatmap showing the clustering of immunoglobulin-related genes in four groups based on log-relative abundances. (G) Anti-tumor specific IgG1, IgG2a, and IgG2b levels in the sera of tumor-bearing mice measured as mean fluorescence intensity (MFI). Each point represents an individual mouse (n=27-29). *p<0.05; **p<0.01; ****p<0.0001.

It is known that Ab isotype can influence anti-tumor immunity (35, 36). Next, we collected sera from tumor-bearing mice at day 14 post tumor cell inoculation and measured B16.F10 tumor-specific antibodies and their isotype by incubating sera with tumor cells followed by Flow cytometry. As shown in figure 5G, anti-tumor IgG1, IgG2a, IgG2b Abs were decreased in the B cell c-Maf KO tumor-bearing mice, suggesting modulatory effect of B cell c-Maf signaling on *in vivo* antibody responses.

### B cell profiling in the peripheral blood of melanoma patients

To explore the clinical significance of study, peripheral blood mononuclear cells (PBMCs) of healthy donors (HD) were stimulated with recombinant human IL-4, CD40L, and 30% of human melanoma A375 conditioned medium (CM). IL-4, CD40L, or A375 CM alone had no effect on c-Maf expression in human B cells whereas combination of IL-4, CD40L and A375 CM significantly upregulated c-Maf expression (figure 6A). Consequently, the combination of IL-4, CD40L and A375 CM significantly upregulated IL-10 production by human B cells, and inhibition of c-Maf using c-Maf inhibitor Nivalenol (NIV) down-regulated IL-10 production by B cells (figure 6B). We further examined human IgG1^+^ B cell differentiation, A375 CM treatment significantly down-regulated IL-4 plus CD40L induced human IgG1^+^ B cell differentiation (online supplemental figure 4A).

**Figure 6.**
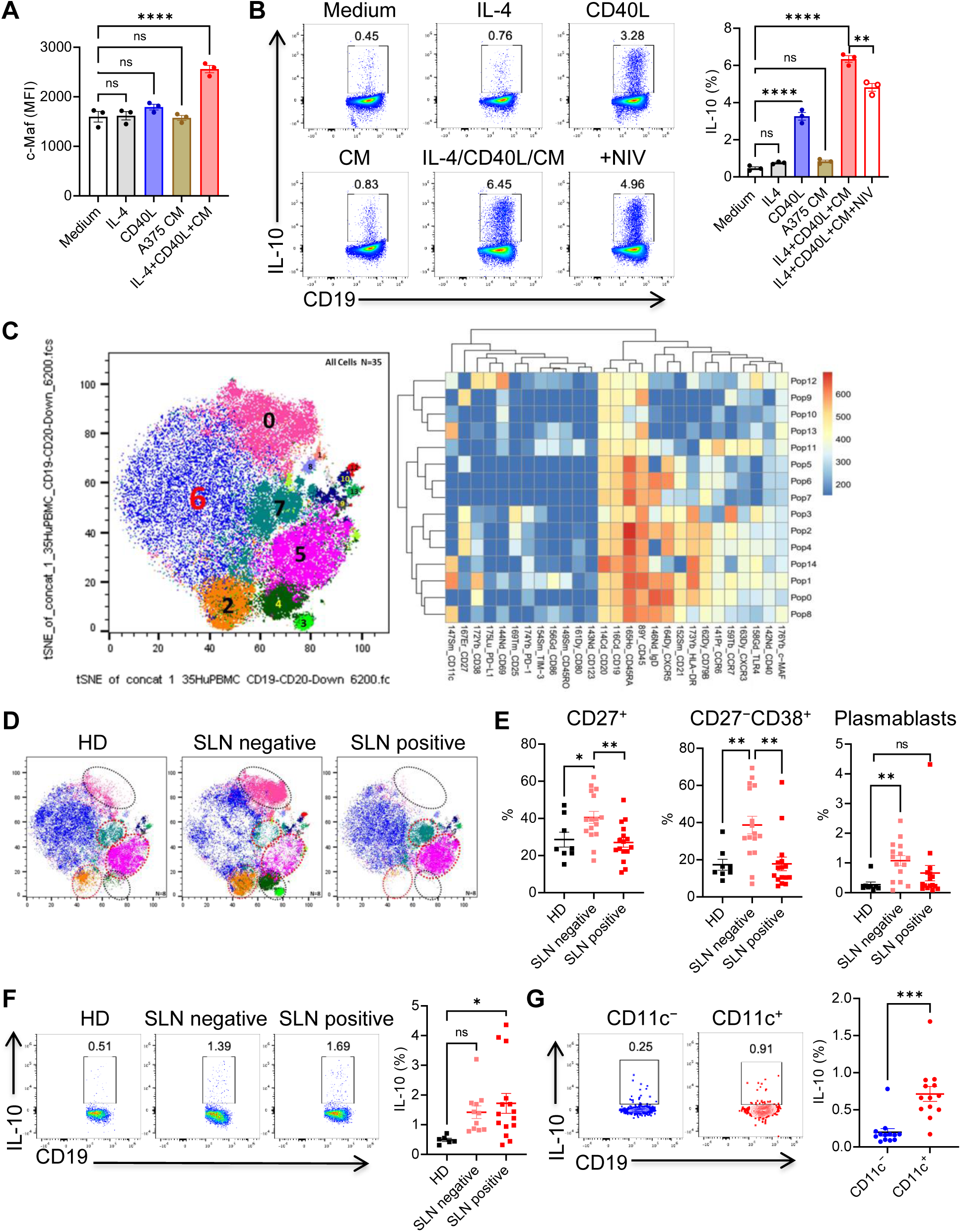
B cell profiling in the peripheral blood of melanoma patients. (A) Mean fluorescence intensity (MFI) of c-Maf expression in human B cells after 3 days culture. (B) PBMCs were stimulated with IL-4/CD40L and A375 conditioned medium with or without c-Maf inhibitor Nivalenol (0.1 μM). IL-10 production by B cells was determined by intracellular cytokine staining. (C) viSNE analysis of CyTOF immunophenotyping of B cells from healthy donors and melanoma patients, all samples combined (n=35, left). FlowSOM clustering into 15 B cell subsets showing as a normalized expression heatmap (right). (D) viSNE analysis of CyTOF immunophenotyping in healthy donors and melanoma patients. (E) Percentages of CD27^+^ B cells, CD27^−^CD38^+^ B cells, and plasmablasts within CD19^+^ B cells. (F) PBMCs were stimulated with IL-4/CD40L for 3 days followed by PMA/Ionomycin/LPS stimulation for 4 hours. IL-10 production by B cells was determined by intracellular cytokine staining. Each point represents an individual person (n=6-16). (G) Freshly thawed PBMCs from melanoma patients were stimulated with PMA/Ionomycin/LPS stimulation for 4 hours. IL-10 production by CD11c^−^ and CD11c^+^ B cells was determined. ns, not significant; *p<0.05; **p<0.01; ***p<0.001; ****p<0.0001; ns, not significant. CM, conditioned medium; HD, healthy donors; SLN, sentinel lymph node. PBMC, peripheral blood mononuclear cells; IL-4, interleukin 4; CD40L, CD40 ligand. NIV, Nivalenol.

Next, we examined B cell profiling and c-Maf expression in the PBMC of healthy donors and melanoma patients with tumor-negative and –positive sentinel lymph nodes (SLN) using CyTOF. FlowSOM analysis identified six major B cell subsets and nine small populations after gating on CD19 and CD20 double-positive B cells. Relatively low c-Maf expression was observed in several subsets, such as population 1, 8, 12, and 13 (figure 6C). CD27 is one of the most used markers to define human memory B cells (37). Three CD27^+^ B cell subsets (population 2, 4, and 3) were accumulated in the peripheral blood of SLN-negative melanoma patients and remarkably reduced in SLN-positive patients. Similarly, SLN-negative melanoma patients had an increased subset of CD27^−^CD38^+^ B cells (population 0), representing a transitional B cell phenotype (38). This subset was decreased in SLN-positive patients compared to that in SLN-negative patients and healthy donors (figure 6D). The changes of CD27^+^ B cells and CD27^−^CD38^+^ B cells were further verified by Flow cytometry analysis of PBMC samples (figure 6E). We also calculated the relative proportion of plasmablasts (CD19^+^CD20^−^CD27^+^CD38^+^) and showed an increase of plasmablasts in SLN negative melanoma patients (figure 6D).

Then, PBMC were cultured with IL-4 and CD40L for 3 days and IL-10 production was examined by intracellular cytokine staining. As shown in figure 6F, IL-10 expression was significantly increased in B cells from melanoma patients, especially SLN-positive patients. To determine the relationship between c-Maf and IL-10 expression in melanoma patients, freshly recovered PBMCs from melanoma patients were restimulated with PMA/Ionomycin/LPS for IL-10 expression. As CD11c is highly expressed in c-Maf positive cluster 1, 8, and 13, we compared IL-10 expression in CD11c^−^ B cells (no c-Maf expression) and CD11c^+^ B cells (c-Maf positive). Significantly higher level of IL-10 was observed in CD11c^+^ B cells compared to CD11c^−^ B cells (figure 6G), suggesting a correlation between c-Maf and IL-10 expression in human B cells. Further, we examined IgG1^+^ B cells and no difference was observed in melanoma patients compared to healthy donors (online supplemental figure 4B).

Lastly, we examined the correlation between c-Maf expression and B cell infiltration by using publicly available web platform TIMER2.0 (http://timer.cistrome.org). As shown in supplemental figure 4C, expression of c-Maf is positively correlation with B cell infiltration and CD19 gene expression in melanoma as well as pancreatic adenocarcinoma (PAAD). JAK3 and STAT6 activation is essential in IL-4 signaling pathway (39). As we observed c-Maf upregulation in presence of IL-4 and CD40L, we further analyzed the correlation between c-Maf^+^ B cells and IL-4 signature in melanoma and pancreatic adenocarcinoma (PAAD), the expression of c-Maf was positively correlated with gene expression of JAK3 and STAT6.

Together, these results demonstrate that substantial changes of memory B cells, transitional B cells, and IL-10-producing B cells are correlated with the stage of melanoma and presence of SLN metastasis.

## Discussion

Emerging data have demonstrated that B cells play important roles in tumor progression. However, the role of B cells in anti-tumor immunity remains controversial. Both pro-tumor and anti-tumor roles of B cells have been reported in the context of tumor immunology (8, 12, 13, 16, 26). The controversy may be due to the heterogeneity in B cell populations as the balance among the subtypes may impact tumor progression (16, 17, 40). Further, tumor type and location are believed to contribute to the complex functions of B cells in anti-tumor immunity.

Therefore, a full understanding of B cells in tumor development is still needed. This study revealed a critical role of c-Maf expression by B cells in promoting tumor growth in two tumor models, PDAC and melanoma.

B cells secrete a very low level of IL-10 at a steady while specific stimulatory signals, such as BCR, CD40, TLR, and cytokine IL-21/IL-6, are involved in the induction of IL-10^+^ B cells. c-Maf is known for the regulation of IL-10 in B cells (22). Additionally, transcriptional factor STAT3 also modulates IL-10 production (41). However, the roles of IL-10 in anti-tumor immunity remain elusive because both pro-tumor and anti-tumor effects have been reported. For example, IL-10 and PD-1 cooperated to limit tumor-specific CD8^+^ T cells (38). Blockade of IL-10 potentiated anti-tumor immune function in human colorectal cancer liver metastases (39). On the other hand, studies demonstrated that IL-10 can induce anti-tumor effect by preventing CD8^+^ T cell apoptosis (40) and reprogramming of exhausted CD8^+^ T cells (41, 42). Further, IL-10^+^ B cells only represent a relatively small fraction of total B cells. Our data revealed that B16.F10 tumor development increases IL-10 production in B cells from TDLN. However, we observed more circulating follicular B cell accumulation in the pancreas upon KPC tumor progression, which led to down regulation of IL-10^+^ B cells. A recent study further reported that loss of B cell specific IL-10 had no effect on B16.F10 melanoma progression (12). Together, these data suggest that IL-10-independent mechanisms may contribute to the anti-tumor effect of B cell c-Maf deficiency. In this study, we defined different anti-tumor mechanisms of B cell c-Maf deficiency in two tumor models. Specifically, c-Maf signaling modulates the pro-inflammatory phenotype and tumor-specific antibody responses in the KPC tumor-bearing pancreas and TDLN of melanoma, respectively.

Previous studies mainly focused on production of immunosuppressive cytokines by a small subset of B cells (14, 15, 17). We are interested in evaluation of the overall phenotype of B cells as a whole population in KPC tumor development. Our data demonstrated that B cells display follicular B cell phenotype (CD23^+^CD21^low^) upon tumor progression, which was associated with an increase of IgD^+^IgM^low^ B cells. Although percentage of IL-10^+^ B cells was decreased, isolated total B cells exhibited a decreased antigen-presenting function. A previous study showed that tumor B cells are proinflammatory with upregulation of several proinflammatory cytokines and chemokines which are critical for the recruitment of T cells, macrophages, and dendritic cells (28). Our RNA-seq analysis and cytokine/chemokine array data further revealed that these pro-inflammatory cytokine and chemokines are upregulated in c-Maf deficient B cells. A recent study reported that the pro-inflammatory chemokines can boost NK and T cell recruitment leading immunological tumor control (33). These data suggest that induction of chemokines through c-Maf inhibition could be a powerful approach to immunologically control PDAC, which is regarded as a “cold” tumor with low T cell and NK cell infiltration in the tumor microenvironment (42, 43). This conclusion is also supported by human studies suggesting that the inflammatory phenotype of B cells is helpful for the anti-tumor response in the tumor microenvironment (44, 45).

Tumor-specific antibodies can either promote tumor growth or exhibit immunostimulatory effects, which may relate to the antibody isotypes (35, 36). Mouse IgG1 (functional equivalent to human IgG4^+^) has been shown to inhibit anti-tumor immunity whereas mouse IgG2a/2b (equivalent to human IgG1^+^) contributes to anti-tumor immunity (3, 35, 46). High IgG/Ig ratio was found to associate with increased survival probability in melanoma (47). Our studies revealed important roles of c-Maf in regulating antibody production. In the mouse B16.F10 melanoma model, knockdown of c-Maf in B cells resulted in down-regulation of genes related to immunoglobulin production and circulating immunoglobulin complex. Serum tumor-specific antibodies, including IgG1 and IgG2a/2b, were decreased in B cell c-Maf KO tumor-bearing mice. In human B cells, tumor-conditioned medium drove down-regulation of human IgG1^+^ B cell differentiation. However, we observed no changes of IgG1^+^ B cell differentiation from PBMC between healthy donors and melanoma patients. Future studies will investigate how tumor-specific antibodies impact tumor progression by studying macrophages in the tumor microenvironment to reveal possible interaction between B cells and macrophages.

Our findings highlight pro-tumor functions of c-Maf^+^ B cells. Hence, approaches to inhibit c-Maf signaling can be translated into patients with pancreatic cancer and melanoma.

Understanding of B cells in anti-tumor immunity and association with the response to immune checkpoint blockade (ICB) therapies remains insufficient with inconsistent data from studies (32, 48). Given the phenotypic and functional heterogeneity of B cells, further investigation of targeting immunosuppressive B cell subset will lead to the improvement of ICB treatment. A limitation of this study is using B16.F10 melanoma model which has very few B cells in the tumor microenvironment. We focus on B cells in TLDN because of their significant expansion and requirement for lymphangiogenesis and melanoma metastasis (49, 50).

## MATERIALS AND METHODS

### Mice and tumor models

Conditional B cell c-Maf knockout (KO) mice were generated by intercrossing c-Maf^f/f^ mice (20) with CD19^Cre^ mice (Jackson Laboratory, strain #: 006785). IL-10^gfp^ reporter mice (Vert-X, stain #: 014530) and OT-II transgenic mice (strain #: 004194) were purchased from Jackson Laboratory. Both male and female mice, 6 to 10 weeks old, were used in studies. PDAC tumor model was established as described in our recent studies (51, 52). 1×10^5^ KPC-GFP cells were mixed with Matrigel (Corning) at 1:1 ratio, and 50 μl mixture was orthotopically implanted into the tail of the pancreas of 8-10 weeks old B cell c-Maf KO (CD19^Cre^-c-Maf^f/f^) and control (CD19^wt^-c-Maf^f/f^) mice. Pancreas were collected at day 18-21 post tumor cell implantation. Pancreas weight and GFP^+^ tumor cells within viable CD45^−^ pancreas cells were used to determine the tumor burden. For melanoma mouse model, 5×10^5^ B16.F10 cells in 100 μl PBS were subcutaneously (s.c.) injected into the flank of B cell c-Maf KO and control mice. Tumors were measured twice a week with a caliper. The tumor volumes were calculated using the formula V = (width×width×length)/2. All animal experiments were approved by the Institutional Animal Care and Use Committee of the University of Louisville (#22115).

### Preparation of single cell suspension and flow cytometry

Pancreas were cut into smaller pieces and digested with digestion buffer comprised of 300 U/ml collagenase I and 80 U/ml DNase (Sigma) in complete RPMI1640 medium for 30 min at 37°C. After incubation, the digestion was immediately stopped by addition of 5 ml cold complete medium. The suspension was then filtered through a 40 µm cell strainer into petri dishes and extra tissue chunks were further mashed with syringe columns. The red blood cells were lysed by adding ACK lysis buffer for about 1 min and washed twice with complete medium. For flow cytometry, the cells were blocked in the presence of anti-CD16/CD32 at 4°C for 10 min and stained on ice with the appropriate antibodies and isotype controls in PBS containing 1% FBS. The fluorochrome-labeled antibodies against mouse CD45, CD19, CD21, CD23, CD5, CD43, IgM, IgD, CD9 and their corresponding isotype controls were purchased from BioLegend.

Fixable Viability Dye eFluor™ 780 was from Thermo Fisher Scientific. The samples were acquired using Cytek Aurora cytometry and analyzed using FlowJo software.

### Intracellular staining for cytokine and c-Maf expression

Cells were stimulated with PMA plus Ionomycin in the presence of protein transport blocker Brefeldin A for 4-6 hours. After staining with surface markers, the cells were then washed, fixed, and permeabilized followed BioLegend’s protocol. Cells were resuspended in the permeabilization buffer and stained with antibodies against mouse IFN-γ and TNF-α, or the respective isotype control overnight at 4 °C. For IL-10 production, cells were stimulated with PMA, Ionomycin, and LPS (5 μg/ml) in the presence of Brefeldin A for 4-6 hours followed by intracellular staining using anti-IL-10 Ab. For c-Maf expression measurement, cells were stained with surface markers followed by fixation and permeabilization using eBioscience Foxp3/Transcription Factor Staining Buffer Set. Cells were then stained with anti-human/mouse c-Maf (clone sym0F1, Invitrogen) overnight at 4 °C. CD9^+^ B cells were further sorted from control and c-Maf KO mice, and c-Maf expression was evaluated by using Western blotting. Antibodies used in flow cytometry and Western blotting were listed in supplementary table 1.

### B cell stimulation, proliferation, antigen presenting function, and immunosuppression

Viable CD45^+^CD19^+^ B cells were isolated from tumor-free and KPC tumor-bearing pancreas or TDLN by using FACS aria III cell sorter. After CFSE labeling, cells were stimulated with LPS (1 μg/ml) for 72 hours. B cell proliferation was determined by measuring CFSE division using Flow cytometry after staining with anti-mouse CD19 Ab. For B cell antigen presentation assay, purified B cells were co-cultured with FACS-isolated CD3^+^CD4^+^ T cells from OT-II transgenic mice in the presence of ovalbumin (OVA, 200 μg/ml) for 4 days. T cell proliferation was evaluated by measuring CFSE division using Flow cytometry. For cytokine and chemokine production, sorted viable CD19^+^ B cells were stimulated with PMA/Ionomycin/LPS (5 μg/ml) for 24 hours. Expression of cytokine/chemokine in the culture supernatants was measured by using mouse cytokine array kit (R&D Systems). In some experiment, splenocytes from IL-10^gfp^ reporter mice were cultured with CD40L– and BAFF-expressing feeder cells (CD40LB) in the presence of IL-4 (10 ng/ml) for 4 days followed by additional 3 days culture in the presence of IL-21 (20 ng/ml) (53). IL-10^+^ and IL-10^−^ B cells were sorted and co-cultured with anti-CD3 mAb-activated CD4^+^ and CD8^+^ T cells for 3 days. The IFN-γ production by T cells was evaluated by intracellular cytokine staining and Flow cytometry.

### Anti-tumor antibody measurement

Mouse sera were isolated from B16.F10 tumor-bearing mice (day 14-16 after tumor cell inoculation) and incubated with B16.F10 tumor cells at 1:1 dilution at 4 °C for 30 minutes. After twice washing with PBS, cells were stained with fluorescein-conjugated anti-mouse IgG1, IgG2a, IgG2b Abs. The anti-tumor antibodies were determined by measuring mean fluorescence intensity (MFI) using Flow cytometry.

### RNA sequencing and analysis

B cells (CD45^+^CD19^+^) were sorted from the pancreas of KPC tumor-bearing mice and TDLN of B16.F10 tumor-bearing mice by using BD FACS Aria III. Fixable viability dye was used to exclude dead cells. A post sort analysis was performed to determine the purity of B cells with approximately 90-95% purity. RNA was extracted using a QIAGEN RNAeasy Kit (QIAGEN). The quantity of the purified RNA samples was measured by the RNA High Sensitivity Kit in the Qubit Fluorometric Quantification system (Thermo Fisher Scientific). Library preparation, sequencing, and analysis were performed by LC Sciences (https://lcsciences.com). RNA-seq data (fastq files) have been deposited at Sequence Read Archive with reference PRJNA1147765.

### Human B cell differentiation

The peripheral blood mononuclear cells (PBMCs) were isolated from the blood of healthy donors using Ficoll–Hypaque centrifugation. Cells were cultured in complete RPMI1640 medium and stimulated with recombinant human CD40 ligand (CD40L, 500 ng/ml, BioLegend), IL-4 (20 ng/ml), and 30% conditioned medium of human melanoma cell line A375. After 3 days culture, cells were harvested for c-Maf expression measurement using intracellular staining. For IL-10 production, cells were restimulated with PMA/Ionomycin/LPS for 4-6 hours in the presence of Brefeldin A. IL-10 production in B cells was measured by using intracellular cytokine staining. In some experiments, cells were cultured in the presence of c-Maf inhibitor Nivalenol (NIV, 100 nM; Cayman Chemical).

### PBMC collection from melanoma patients

8-10 ml of peripheral blood was collected from treatment-naïve melanoma patients including both tumor-negative and –positive sentinel lymph node (SLN) at the Department of Surgery, University of Louisville. SLN-positive or SLN-negative was defined as evidence, or no evidence of metastatic tumor cells identified by standard hematoxylin and eosin (H&E) staining and immunohistochemistry (IHC). All samples were anonymously coded in accordance with UofL IRB guidelines, and written informed consent was obtained. The PBMCs were isolated from blood and frozen at –140 °C until subsequent analysis. This study was approved by the institutional review boards (IRB) of University of Louisville (#08.0491).

### CyTOF mass cytometry data acquisition and data analysis

Cells from TDLN of B16.F10 tumor-bearing mice were stimulated with PMA, Ionomycin, and LPS in the presence of Brefeldin A for 4-6 hours at 37 °C. Cell permeabilization and staining were performed according to the protocol from Fluidigm. Prior to acquisition, cells were suspended in a 1:9 solution of cell acquisition solution: EQ 4 element calibration beads (Fluidigm) at an appropriate concentration at no more than 600 events per second. Data acquisition was performed on the CyTOF Helios system (Fluidigm). FCS files were normalized with Helios instrument work platform (FCS Processing) based on the calibration bead signal used to correct any variation in detector sensitivity. CyTOF data analysis was performed with FlowJo software. Total CD45^+^ cells were gated after removing beads, doublets, and dead cells. tSNE and FlowSOM clustering analysis for CyTOF data were performed using FlowJo Plugins platform. tSNE analysis was performed on all samples combined. Different immune populations were defined by the expression of specific surface and intracellular markers.

For human B cell profiling, PBMCs from healthy donors (n=8) and melanoma patients with tumor-negative SLN (n=16) and –positive SLN (n=16) were used for CyTOF analysis. Cell permeabilization and staining were performed according to the protocol from Fluidigm. Total B cells were gated on CD19 and CD20 double positive cells after removing beads, doublets, and dead cells followed by tSNE and FlowSOM clustering analysis. Antibodies used for CyTOF were listed in supplementary table 2 and 3. The CyTOF (fcs files) have been uploaded to Figshare (https://figshare.com/authors/Chuanlin_DING/19321330).

### Statistical analysis

Data were analyzed using GraphPad Prism software (GraphPad Software). An unpaired Student *t* test and one-way or two-way ANOVA were used to calculate significance, as appropriate. All graph bars are expressed as mean ± SEM. Significance was assumed to be reached at *p < 0.05*. The *p* values were presented as follows: *\*p<0.05, **p<0.01, ***p<0.001,* and *\*\*\*\*p<0.0001*.

## Ethics approval

All animal experiments were approved by the Institutional Animal Care and Use Committee of the University of Louisville (#22115).

## Consent for publication

All blood samples were anonymously coded in accordance with UofL IRB guidelines, and written informed consent was obtained. This study was approved by the institutional review boards (IRB) of University of Louisville (#08.0491).

## Data availability statement

The RNA-seq data (fastq files) have been deposited at Sequence Read Archive with reference PRJNA1147765. The CyTOF data (fcs files) have been uploaded to Figshare (https://figshare.com/authors/Chuanlin_DING/19321330). All other data are available upon reasonable request.

## Contributors

QZ conducted experiment, analyzed data, prepared figures, and wrote the manuscript. HH provided PBMCs of melanoma patients. SL helped with some in vivo experiment involving KPC tumor cell injection and pancreas tissue processing. HL analyzed CyTOF data. YN and XL helped with PBMC processing and B cell c-Maf knockdown validation. KM and JY provided advice on research design, IRB and IACUC protocol approval, and performed a critical review. CD analyzed data, prepared figures, wrote the manuscript, and is responsible for the overall content as the guarantors.

## Acknowledgements

CyTOF data acquisition and analysis were performed at UofL BCC Functional Immunomics Core supported by NIH P20GM135004.

