## Supplemental material and data for "B cell c-Maf signaling promotes tumor progression in animal models of pancreatic cancer and melanoma"

**Supplemental Table 1.** Antibodies used in Flow cytometry and Western blot

| Antibodies | Company and catalog number |
|---|---|
| PerCP/Cyanine5.5 anti-mouse CD45 (clone S18009F) | BioLegend Cat # 157208 |
| PE/Cyanine7 anti-mouse CD45 (clone S18009F) | BioLegend Cat # 157206 |
| PerCP/Cyanine5.5 anti-mouse IgM (clone RMM-1) | BioLegend Cat # 406512 |
| FITC anti-mouse CD21/CD35 (clone 7E9) | BioLegend Cat # 123408 |
| APC anti-mouse CD19 (clone 6D5) | BioLegend Cat # 115512 |
| PE anti-mouse IgM (clone RMM-1) | BioLegend Cat # 406508 |
| PerCP/Cyanine5.5 anti-mouse IgD (clone 11-26c.2a) | BioLegend Cat # 405709 |
| PE/Cyanine7 anti-mouse CD5 (clone 53-7.3) | BioLegend Cat # 100621 |
| FITC Anti-Mouse CD43 (clone S7) | BD Bioscience Cat # 553270 |
| APC anti-mouse CD4 (clone GK1.5) | BioLegend Cat # 100412 |
| FITC anti-mouse CD8a (clone 53-6.7) | BioLegend Cat # 100706 |
| APC anti-mouse CD19 (clone 6D5) | BioLegend Cat # 115512 |
| PE anti-mouse IL-10 (clone JES5-16E3) | BioLegend Cat # 505007 |
| PE anti-mouse IFN-γ (clone XMG1.2) | BioLegend Cat # 505808 |
| PE anti-mouse IgG1 (clone RMG1-1) | BioLegend Cat # 406608 |
| PE anti-mouse IgG2a (clone RMG2a-62) | BioLegend Cat # 407108 |
| PE anti-mouse IgG2b (clone RMG2b-1) | BioLegend Cat # 406708 |
| PE anti-human CD19 (clone HIB19) | BioLegend Cat # 302208 |
| PerCP/Cyanine5.5 anti-human IgD (clone IA6-2) | BioLegend Cat # 348208 |
| PE/Cyanine7 anti-mouse/rat/human CD27 (clone LG.3A10) | BioLegend Cat # 124215 |
| APC anti-human IL-10 (clone JES3-19F1) | BioLegend Cat # 506807 |
| Alexa Fluor 700 anti-human CD11c (clone Bu15) | BioLegend Cat # 337220 |
| PerCP-eFluor™ 710 c-MAF monoclonal antibody (sym0F1) | Invitrogen Cat # 46-9855-42 |
| PE c-MAF monoclonal antibody (sym0F1) | Invitrogen Cat # 12-9855-42 |
| c-Maf Antibody (M-153) for Western blotting | Santa Cruz Cat # sc-7866 |
| Fixable Viability Dye eFluor™ 780 | Thermo Fisher Scientific Cat # 65-0865-14 |

**Supplemental Table 2.** Anti-mouse antibodies for CyTOF

| | Tag | Target | Clone | Company and catalog number |
|---|---|---|---|---|
| 1 | 89Y | CD45 | 30-F11 | Fluidigm Cat # 3089005B |
| 2 | 141Pr | TNF-α | MP6-XT22 | Fluidigm Cat # 3141013B |
| 3 | 142Nd | CD11c | N418 | Fluidigm Cat # 3142003B |
| 4 | 143Nd | CD69 | H1.2F3 | Fluidigm Cat # 3143004B |
| 5 | 144Nd | IL-2 | JES6-5H4 | Fluidigm Cat # 3144002B |
| 6 | 145Nd | CD4 | RM4-5 | Fluidigm Cat # 3145002B |
| 7 | 146Nd | F4/80 | BM8 | Fluidigm Cat # 3146008B |
| 8 | 148Nd | CD103 | 2E7 | BioLegend Cat # 121402 |
| 9 | 149Sm | CD19 | 6D5 | Fluidigm Cat # 3149002B |
| 10 | 150Nd | Ly-6C | NK1.4 | Fluidigm Cat # 3150010B |
| 11 | 151Eu | CD25 | 3C7 | Fluidigm Cat # 3151007B |
| 12 | 152Sm | CD3e | 145-2C11 | Fluidigm Cat # 3152004B |
| 13 | 153Eu | CD274/PD-L1 | MIH5 | Fluidigm Cat # 3153031B |
| 14 | 155Gd | IL-10 | JES5-16E3 | BioLegend Cat # 505029 |
| 15 | 156Gd | CCR2 | 475301R | R&D System Cat # MAB55381R |
| 16 | 158Gd | Foxp3 | FJK-16s | Fluidigm Cat # 3158003A |
| 17 | 159Tb | PD-1 | 29F.1A12 | Fluidigm Cat # 3159024B |
| 18 | 160Gd | CD62L | MEL-14 | Fluidigm Cat # 3160008B |
| 19 | 161Dy | iNOS | CXNFT | Fluidigm Cat # 3161011B |
| 20 | 162Dy | CD44 | IM7 | Fluidigm Cat # 3162030B |
| 21 | 164Dy | CX3CR1 | SA011F11 | Fluidigm Cat # 3164023B |
| 22 | 165Ho | IFN-γ | XMG1.2 | Fluidigm Cat # 3165003B |
| 23 | 167Er | IL-6 | MP5-20F3 | Fluidigm Cat # 3167003B |
| 24 | 168Er | CD8a | 53-6.7 | Fluidigm Cat # 3168003B |
| 25 | 169Tm | CD206 | C068C2 | Fluidigm Cat # 3169021B |
| 26 | 170Er | NK1.1 | PK136 | Fluidigm Cat # 3170002B |
| 27 | 172Yb | CD11b | M1/70 | Fluidigm Cat # 3172012B |
| 28 | 174Yb | CD223/LAG3 | C9B7W | Fluidigm Cat # 3174019B |
| 29 | 175Yb | CD127/IL7Ra | A7R34 | Fluidigm Cat # 3175006B |
| 30 | 176Yb | CD45R/B220 | RA3-6B2 | Fluidigm Cat # 3176002B |
| 31 | 209Bi | I-A/I-E | M5/114.15.2 | Fluidigm Cat # 3209006B |

**Supplemental Table 3.** Anti-human antibodies for CyTOF

| | **Tag** | **Target** | **Clone** | **Company and catalog number** |
|---|---|---|---|---|
| 1 | 89Y | CD45 | HI30 | Fluidigm Cat # 3089003B |
| 2 | 114Cd | CD20 | 2H7 | BioLegend Cat # 302343 |
| 3 | 116Cd | CD19 | HIB19 | BioLegend Cat # 302247 |
| 4 | 141Pr | CD196/CCR6 | G034E4(11A9) | Fluidigm Cat # 3141003A |
| 5 | 142Nd | CD40 | 5C3 | Fluidigm Cat # 3142010B |
| 6 | 143Nd | CD123 | 6H6 | Fluidigm Cat # 3143014B |
| 7 | 144Nd | CD69 | FN50 | Fluidigm Cat # 3144018B |
| 8 | 146Nd | IgD | IA6-2 | Fluidigm Cat # 3146005B |
| 9 | 147Sm | CD11c | Bu15 | Fluidigm Cat # 3147008B |
| 10 | 149Sm | CD45RO | UCHL1 | Fluidigm Cat # 3149001B |
| 11 | 152Sm | CD21 | BL13 | Fluidigm Cat # 3152010B |
| 12 | 154Sm | TIM-3 | F38-2E2 | Fluidigm Cat # 3154010B |
| 13 | 156Gd | CD86 | IT2.2 | Fluidigm Cat # 3156008B |
| 14 | 158Gd | CD284 | HTA125 | Fluidigm Cat # 3158024B |
| 15 | 159Tb | CD197/CCR7 | G043H7 | Fluidigm Cat # 3159003A |
| 16 | 161Dy | CD80 | 2D10.4 | Fluidigm Cat # 3161023B |
| 17 | 162Dy | CD79b | CB3-1 | Fluidigm Cat # 3162008B |
| 18 | 163Dy | CXCR3 | G025H7 | Fluidigm Cat # 3163004B |
| 19 | 164Dy | CXCR5 | RF8B2 | Fluidigm Cat # 3164029B |
| 20 | 165Ho | CD45RA | HI100 | BioLegend Cat # 304143 |
| 21 | 167Er | CD27 | L128 | Fluidigm Cat # 3167006B |
| 22 | 169Tm | CD25 | 2A3 | Fluidigm Cat # 3169003B |
| 23 | 172Yb | CD38 | HIT2 | Fluidigm Cat # 3172007B |
| 24 | 173Yb | HLA-Dr | L243 | Fluidigm Cat # 3173005B |
| 25 | 174Yb | CD279/PD1 | EH12.2H7 | Fluidigm Cat # 3174020B |
| 26 | 175Lu | CD274/PD-L1 | 29E.2A3 | Fluidigm Cat # 3175017B |
| 27 | 176Yb | c-Maf | Polyclone | Thermo Fisher Cat # PA5-23179 |

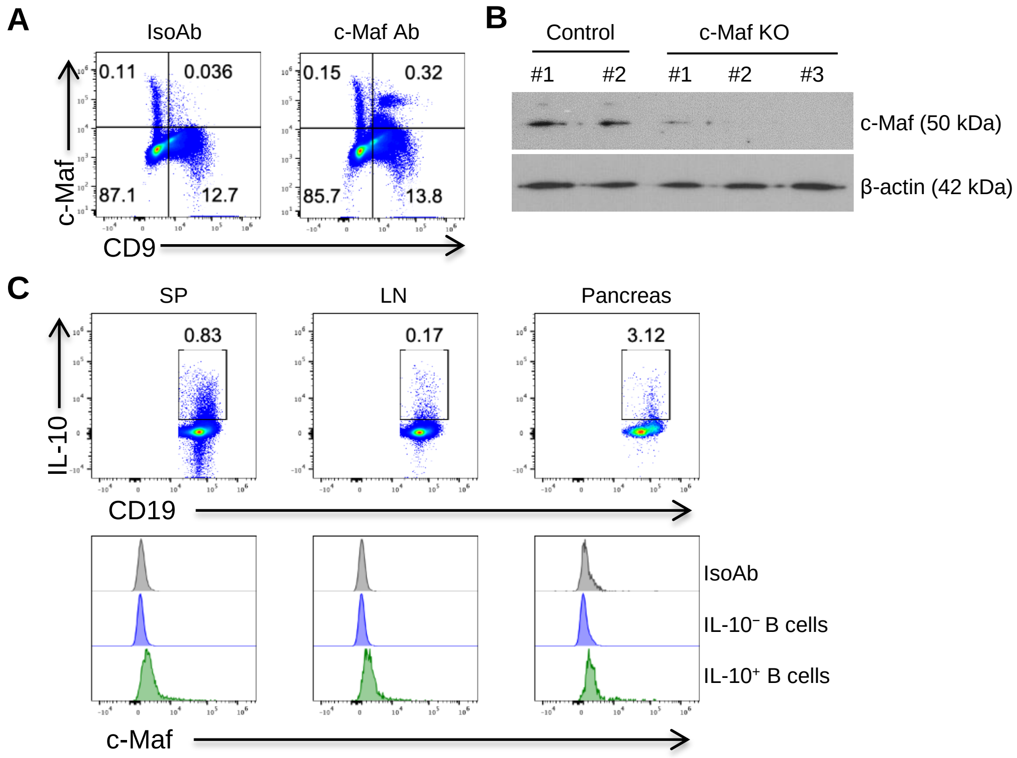


**Supplemental Figure 1.** (A) Splenocytes were stained with CD19 and CD9 antibodies followed by intracellular staining with isotype antibody and c-Maf antibody. Cells were gated on viable $CD19^+$ cells. (B) $CD9^+$ B cells were sorted from control (n=2) and conditional B cell c-Maf KO mice (n=3). c-Maf expression was determined by using Western blotting. (C) Spleen, lymph node, and pancreas were collected from naïve IL-$10^{gfp}$ reporter mice, and IL-10 expression was determined by measuring GFP fluorescence. c-Maf expression in naïve IL-$10^-$ and IL-$10^+$ B cells was determined by intracellular staining and Flow cytometry.

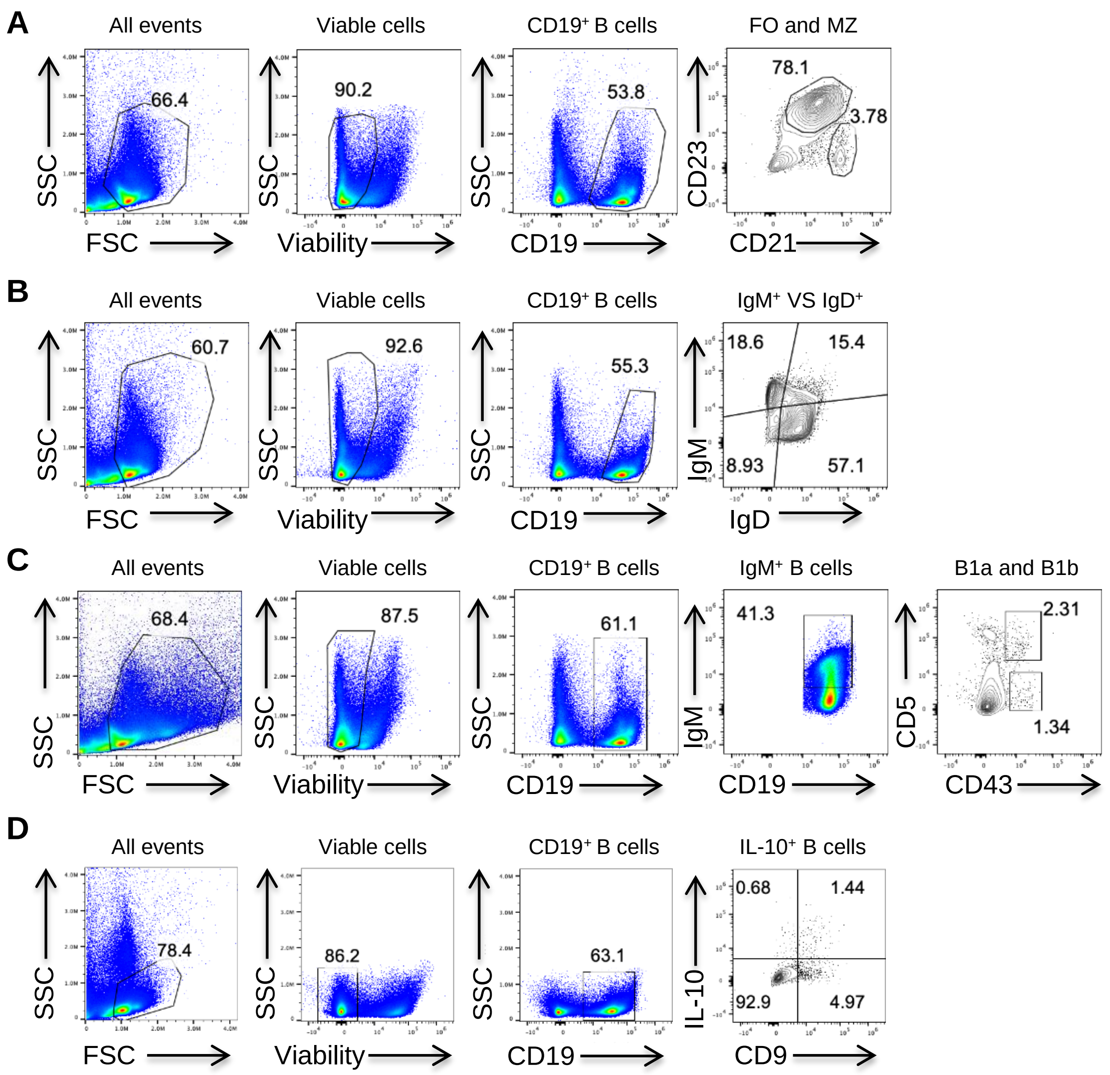


**Supplemental Figure 2.** Gating strategy used (A) for identification of follicular B cells (FO) and marginal zone (MZ) B cells; (B) for identification of IgM$^{+}$ and IgD$^{+}$ B cells; (C) for identification of B1a and B1b; (D) for identification of CD9$^{+}$IL-10$^{+}$ B cells in figure 1.

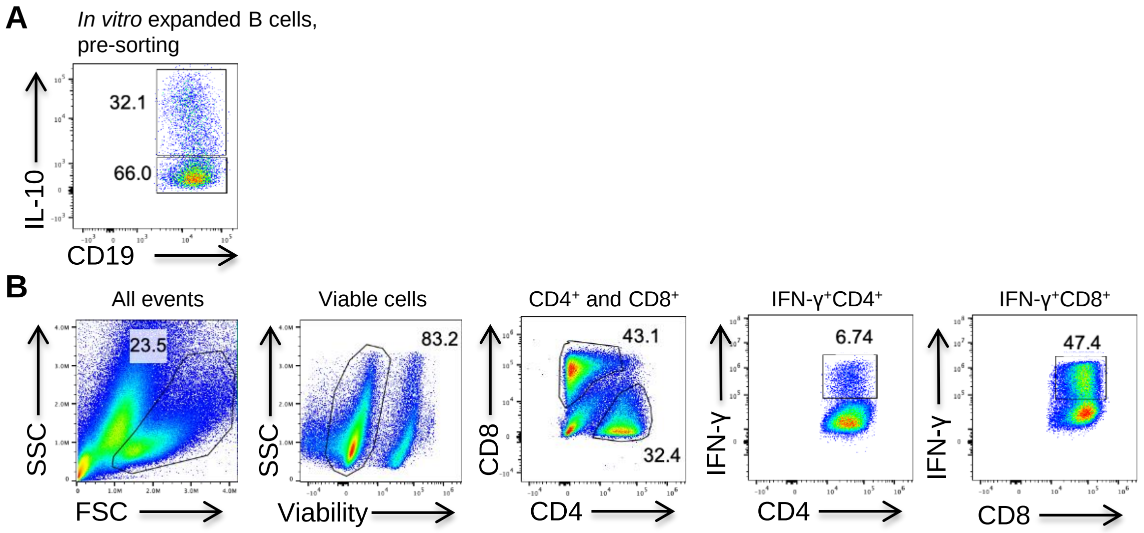


**Supplemental Figure 3.** Gating strategy used for IL-10$^+$ B cell sorting and coculture with CD4, CD8 T cells. (A) Splenocytes of IL-10$^{gfp}$ reporter mice were cultured with CD40L- and BAFF-expressing feeder cells (CD40LB) in the presence of IL-4 for 4 days followed by additional 3 days culture in the presence of IL-21. IL-10 positive verse negative was determined by measuring GFP fluorescence in the *in vitro* expanded B cells. (B) Gating strategy of co-culture of B cells and T cells. IL-10$^+$ and IL-10$^-$ B cells were sorted and cocultured with anti-CD3 mAb activated CD4$^+$ and CD8$^+$ T cells for 3 days. The IFN-γ production by CD4$^+$ and CD8$^+$ T cells was evaluated by intracellular cytokine staining and Flow cytometry.

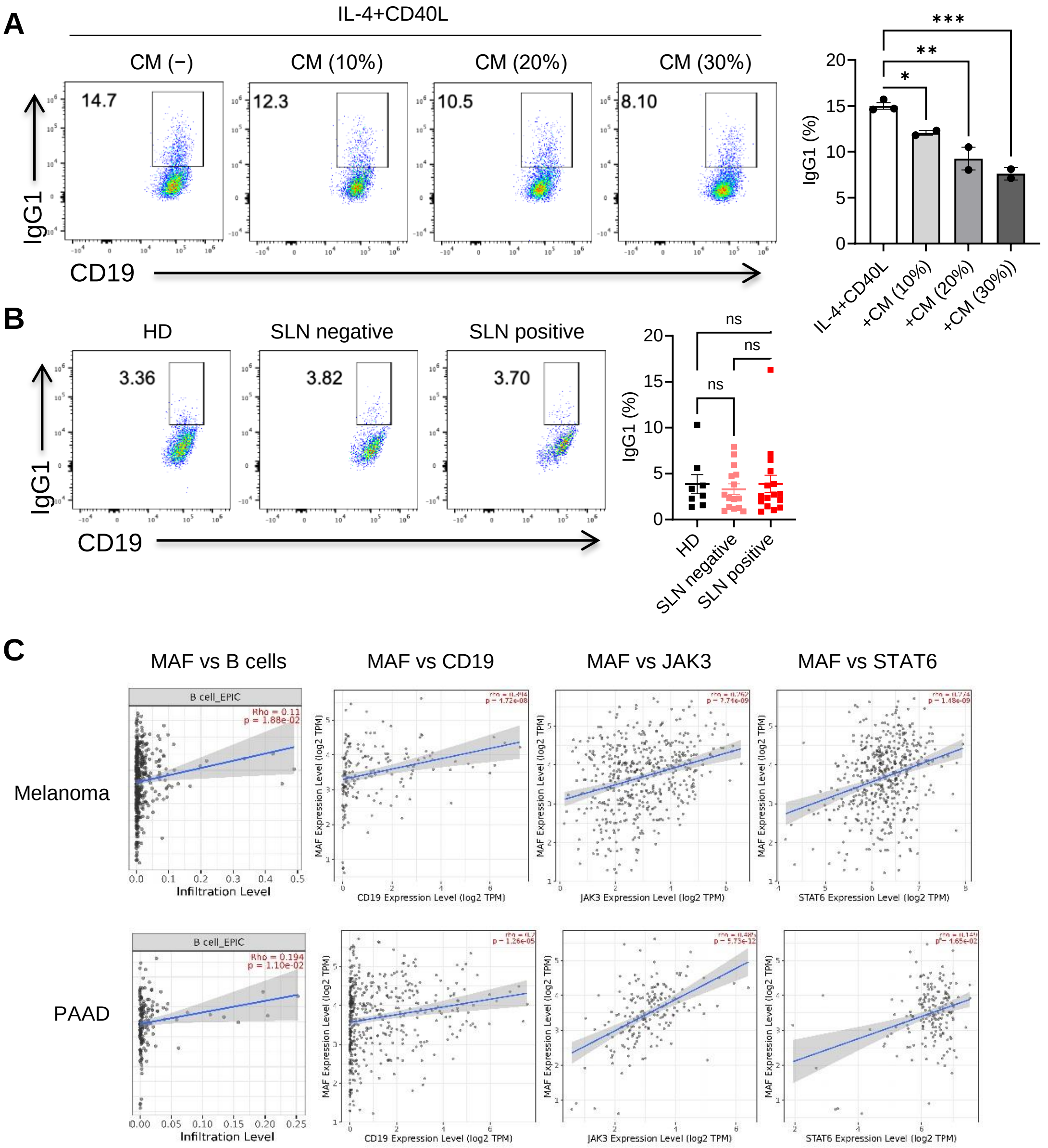


**Supplemental Figure 4.** (A) PBMCs from healthy donors were cultured in the presence of IL-4 and recombinant CD40L with or without A375 conditioned medium for 3 days. IgG1 positive B cells were determined by using Flow cytometry. (B) PBMCs from healthy donors, SLN negative and SLN positive melanoma patients were cultured in the presence of IL-4 and recombinant CD40L for 3 days. IgG1 positive B cells were determined by using Flow cytometry. Summarized data were shown. Each dot represents one person. *$p$ <0.05, **$p$ <0.01, ***$p$ <0.001 by Ordinary one-way ANOVA test. (C) The correlation of MAF and B cell infiltration as well as CD19, JAK3, STAT6 gene expression in melanoma and PAAD patients from The Cancer Genome Atlas (TCGA) using publicly available web platform TIMER2.0.
